# Geographic Genetic Variation in the Introduced Insect *Hyphantria cunea* Drury in Japan Revealed by RAPD Analysis

**DOI:** 10.1101/2025.07.10.664082

**Authors:** Haruhisa Kawasaki, Yasuyuki Shirota

## Abstract

The fall webworm *Hyphantria cunea* Drury (Lepidoptera: Arctiidae) is an invasive pest in Japan that has expanded its range since its accidental introduction in 1945. To investigate the genetic structure and potential source of a northern population recently established in Hirosaki (Aomori Prefecture), we analyzed five geographic populations using the random amplified polymorphic DNA (RAPD) method. Two primers revealed 18 polymorphic bands among 155 individuals collected across five prefectures. Cluster analysis based on Nei’s genetic distances grouped the populations into two major clusters corresponding to the Pacific coast and Sea of Japan coast, with the Hirosaki population showing close affinity to the Pacific coast group. These results suggest possible evolutionary divergence and local adaptation following invasion, and they highlight the utility of RAPD markers in assessing geographic genetic variation in introduced insect populations.

## Introduction

The fall webworm, *Hyphantria cunea* Drury, is a polyphagous defoliator of numerous ornamental and fruit-bearing trees, including cherry, mulberry, plane, and persimmon (Warren and Tadic 1970). Originally native to North America, *H. cunea* was introduced to Japan around 1945 (Umeya and Ito 1977), and has since spread throughout most of the country (Hasegawa 1966; Masaki 1975; Gomi 1996). The species’ initial expansion was studied extensively, and a northern distribution limit was proposed based on thermal requirements (Ito et al. 1968; Masaki 1977), with 1600 day-degree as the survival threshold.

Unexpectedly, since the 1990s, the species has been found beyond this thermal boundary, including in Hirosaki City (40.4°N), Aomori Prefecture. This area, considered climatically marginal for *H. cunea*, supports a bivoltine population that overwinters as pupae. The persistence of this population raises questions about its origin and adaptation. While earlier dispersal was linked to human activity (e.g., transport via roads and railways), natural dispersal via flight is also plausible.

To elucidate the origin of the Hirosaki population and assess the degree of genetic differentiation among regional populations, we employed RAPD analysis. This method allows the detection of polymorphic loci without prior genomic information, making it suitable for evaluating consanguinity among geographically separated populations.

## Materials and Methods

### Sample Collection

Samples of *H. cunea* moths were collected from five prefectures in the Tohoku region of Japan (Aomori, Akita, Iwate, Yamagata, and Miyagi) between June 1992 and October 1993. Collection sites and sample sizes are detailed in Table 1 (see Appendix).

**Table 1.** Sampling locations and geographic coordinates of the five *Hyphantria cunea* populations used in this study. N indicates the number of individuals analyzed per population.

| Population | Location | N | Latitude | Longitude |
| --- | --- | --- | --- | --- |
| AOMORI | Hirosaki City | 59 | 40.4°N | 140.3°E |
| AKITA | Akita City | 27 | 39.4°N | 139.8°E |
| IWATE | Morioka City | 35 | 39.4°N | 141.1°E |
| YAMAGATA | Sakata City | 19 | 38.6°N | 139.5°E |
| MIYAGI | Sendai City | 15 | 38.2°N | 140.4°E |

### DNA Extraction

Genomic DNA was extracted from individual pupae using a modified protocol based on Hills and Moritz (1990). Each sample was homogenized in lysis buffer (10 mM NaCl, 25 mM EDTA, 10 mM Tris-HCl, pH 8.0, and 0.5% SDS). After digestion with proteinase K (0.1 mg/ml) at 37°C for 30 minutes, samples were purified via phenol-chloroform extraction. DNA was precipitated with sodium acetate and ethanol, washed with 70% ethanol, air-dried, and resuspended in TE buffer.

### RAPD-PCR Amplification

RAPD-PCR was performed using two 10-mer primers (OPT-01: 5’-GGGCCACTCA-3’ and OPT-02: 5’-GGAGAGACTC-3’) selected from a screening panel (Operon Technologies). Reactions (25 μl) included 1.25 mM dNTPs, 10× Tth polymerase buffer, 0.5 U Tth polymerase, 5 pmol of each primer, and 1 μl genomic DNA. Amplification was carried out using a Perkin Elmer Thermal Cycler under the following conditions: 5 min at 94°C; 45 cycles of 1 min at 93°C, 1 min at 37°C, and 2 min at 72°C; with a final extension at 72°C for 5 min.

PCR products were electrophoresed on 5% polyacrylamide gels in TBE buffer and stained with ethidium bromide for visualization under UV light.

### Data Analysis

A total of 18 polymorphic RAPD bands ranging from 297 to 1050 bp were scored as present (1) or absent (0). Bands were assumed to represent dominant alleles from homologous loci under Hardy-Weinberg equilibrium. Genetic diversity statistics, including total diversity (HT), within-population diversity (HS), Nei’s gene diversity (H), genetic differentiation (GST), and Nei’s genetic distance (D), were calculated using PopGene 1.32 (Yeh et al. 1999). Cluster analysis was performed using the UPGMA method.

## Results

Eighteen polymorphic RAPD bands were consistently amplified across 155 individuals. The HT values ranged from 0.0142 to 0.5000 depending on the marker, while HS ranged from 0.0141 to 0.4673. The mean GST was 0.1087, indicating moderate genetic differentiation among populations (Table 2).

**Table 2.**
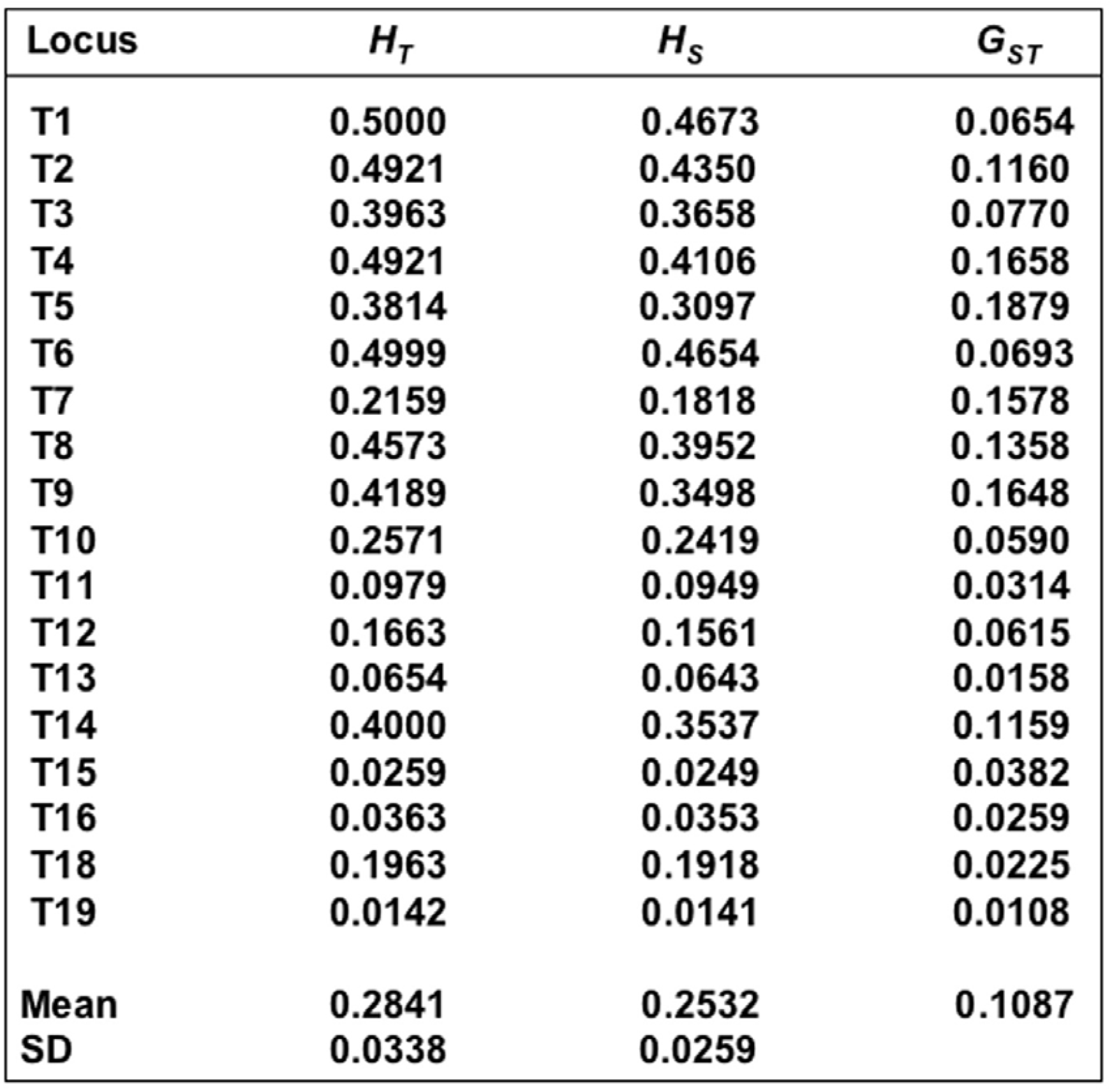
Genetic diversity statistics at 18 RAPD loci among five *Hyphantria cunea* populations. For each marker, total genetic diversity (HT), intra-population diversity (HS), and the coefficient of genetic differentiation (GST) were estimated. Values represent single-locus diversity based on dominant allele presence/absence scoring.

| **Locus** | **$H_T$** | **$H_S$** | **$G_{ST}$** |
| --- | --- | --- | --- |
| **T1** | **0.5000** | **0.4673** | **0.0654** |
| **T2** | **0.4921** | **0.4350** | **0.1160** |
| **T3** | **0.3963** | **0.3658** | **0.0770** |
| **T4** | **0.4921** | **0.4106** | **0.1658** |
| **T5** | **0.3814** | **0.3097** | **0.1879** |
| **T6** | **0.4999** | **0.4654** | **0.0693** |
| **T7** | **0.2159** | **0.1818** | **0.1578** |
| **T8** | **0.4573** | **0.3952** | **0.1358** |
| **T9** | **0.4189** | **0.3498** | **0.1648** |
| **T10** | **0.2571** | **0.2419** | **0.0590** |
| **T11** | **0.0979** | **0.0949** | **0.0314** |
| **T12** | **0.1663** | **0.1561** | **0.0615** |
| **T13** | **0.0654** | **0.0643** | **0.0158** |
| **T14** | **0.4000** | **0.3537** | **0.1159** |
| **T15** | **0.0259** | **0.0249** | **0.0382** |
| **T16** | **0.0363** | **0.0353** | **0.0259** |
| **T18** | **0.1963** | **0.1918** | **0.0225** |
| **T19** | **0.0142** | **0.0141** | **0.0108** |
| **Mean** | **0.2841** | **0.2532** | **0.1087** |
| **SD** | **0.0338** | **0.0259** |  |

Nei’s genetic distances between population pairs ranged from 0.0241 to 0.0729 (Table 3). The lowest genetic distance was observed between Aomori and Iwate populations (D = 0.0241), while Miyagi showed the greatest differentiation from other populations. Although the UPGMA dendrogram (Figure 2) grouped the Aomori population with those on the Sea of Japan coast, a direct comparison of pairwise genetic distances (Figure 3) revealed that Aomori is genetically closest to Iwate, suggesting a more complex pattern of relatedness.

**Table 3.** Pairwise geographic distances (in kilometers; above the diagonal) and Nei’s genetic distances (below the diagonal) among five *Hyphantria cunea* populations in northern Japan. Genetic distances were calculated based on 18 polymorphic RAPD markers.

| Population | Population |  |  |  |  |
| --- | --- | --- | --- | --- | --- |
|  | AOMORI | AKITA | IWATE | YAMAGATA | MIYAGI |
| AOMORI | **** | 113 | 123 | 208 | 274 |
| AKITA | 0.0344 | **** | 94 | 94 | 179 |
| IWATE | 0.0241 | 0.0314 | **** | 151 | 170 |
| YAMAGATA | 0.0383 | 0.0244 | 0.0597 | **** | 121 |
| MIYAGI | 0.0566 | 0.0622 | 0.0729 | 0.0535 | **** |

**Figure 1.**
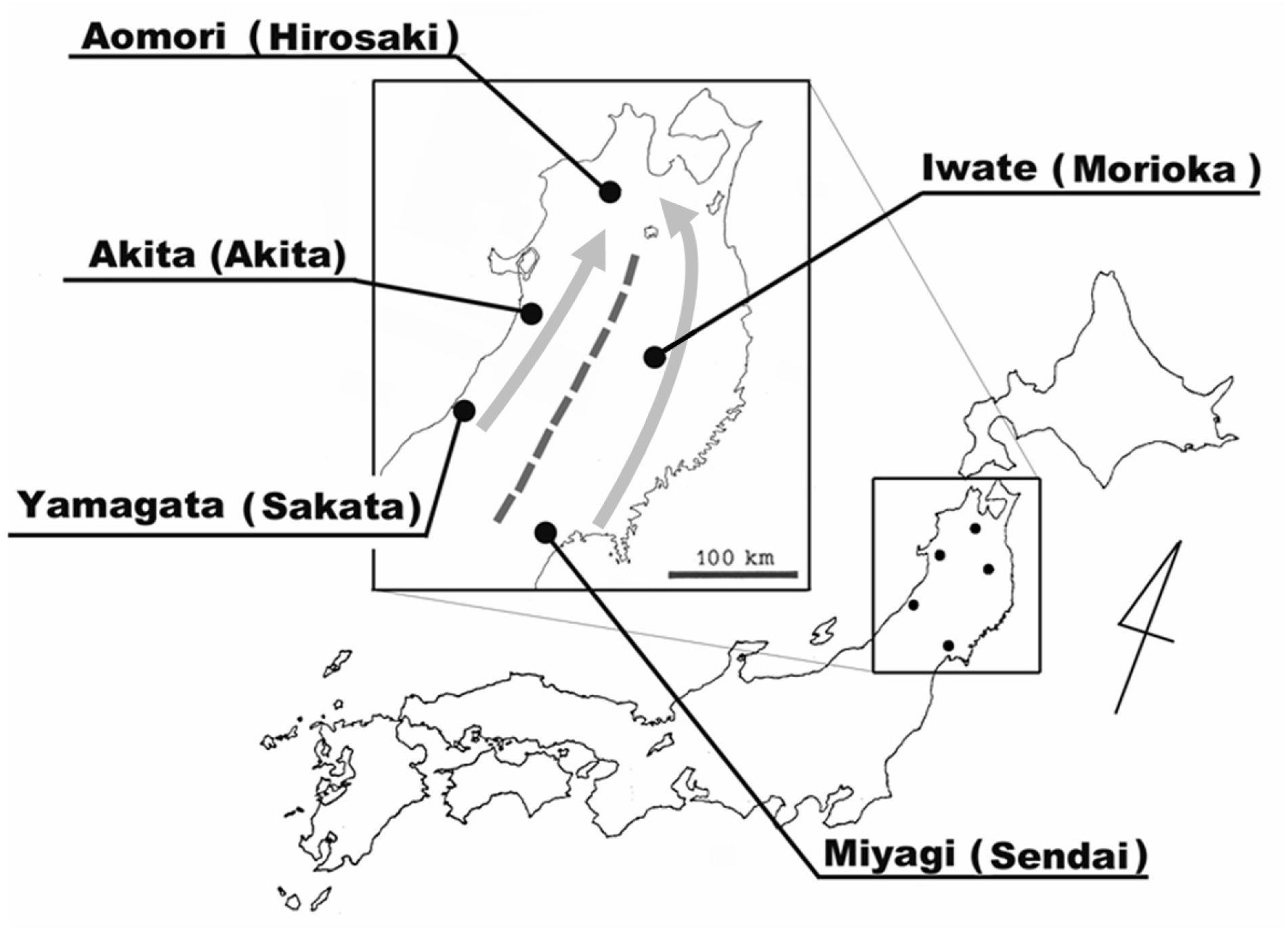
Geographic locations of the five *Hyphantria cunea* collection sites in the Tohoku region of Japan (see also Table 1). Solid dots indicate sampling locations: Hirosaki (Aomori), Morioka (Iwate), Akita (Akita), Sakata (Yamagata), and Sendai (Miyagi). The gray arrows represent hypothesized invasion routes of *H. cunea* into Aomori Prefecture. The dashed line marks the central mountain range that may serve as a geographic barrier between eastern and western populations.

**Figure 2.**
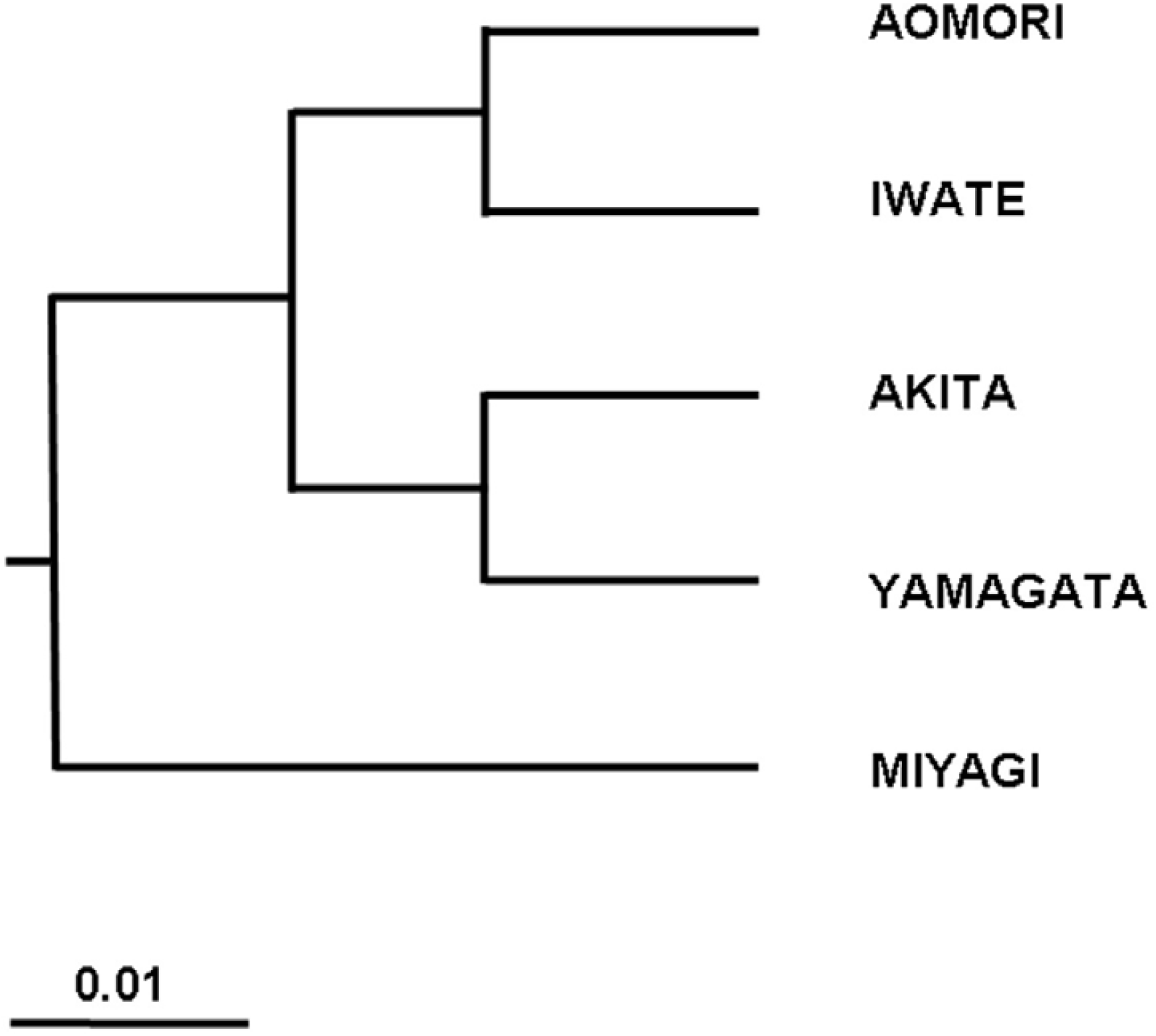
Dendrogram illustrating genetic relationships among five *Hyphantria cunea* populations constructed using the unweighted pair-group method with arithmetic mean (UPGMA) based on Nei’s genetic distance. Clustering reflects geographic and genetic differentiation inferred from RAPD data.

**Figure 3.**
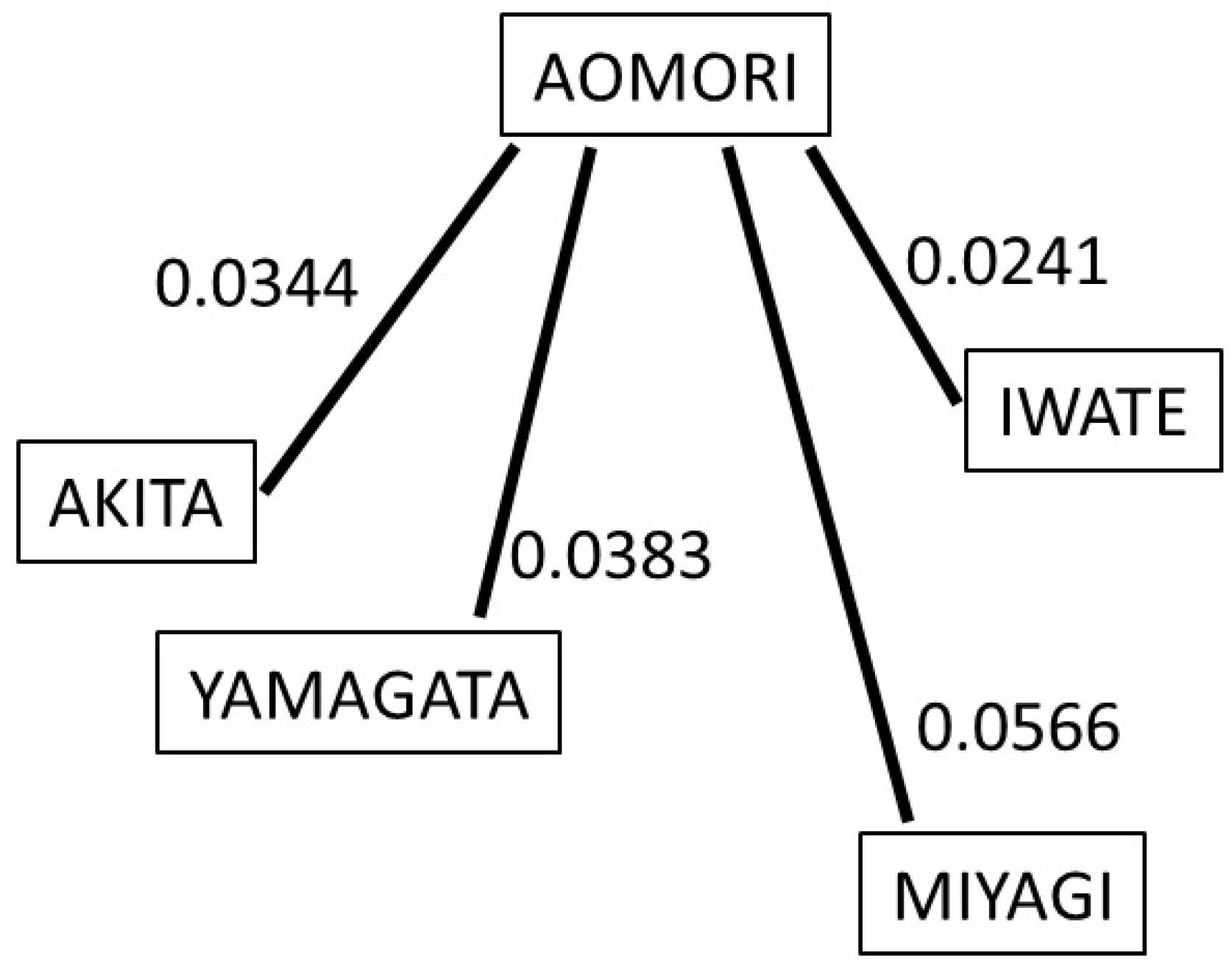
Genetic distances between the Aomori population and each of the other four *Hyphantria cunea* populations, based on pairwise Nei’s genetic distance values derived from RAPD data (see also Table 3). While Aomori was grouped with the Sea of Japan coastal populations in the UPGMA analysis (Figure 2), this diagram highlights its close genetic similarity with Iwate, a Pacific coastal population.

## Discussion

Our results reveal significant genetic structure among *H. cunea* populations in northeastern Japan. Although population-specific bands were not observed, the distribution of polymorphic markers indicates partial genetic isolation. The observed clustering is broadly consistent with geographic location; however, the inclusion of Aomori within the Sea of Japan cluster in Figure 2 contrasts with its close genetic proximity to Iwate in Figure 3.

We propose that the mountainous backbone of the Tohoku region serves as a significant ecological barrier, effectively segregating east and west coasts. Since *H. cunea* cannot readily inhabit mountain forests where natural enemies are abundant, the species likely dispersed northward along lowland corridors on both coasts. The two invasion fronts may have met in the northernmost lowland area— the Tsugaru Plain of Aomori—resulting in a mixing or secondary contact zone.

This hypothesis is supported by the genetic distances depicted in Figure 3, which suggest that the Aomori population retains genetic signals from both east and west coast lineages. While the UPGMA method is useful for summarizing overall trends, pairwise genetic distances offer a more nuanced view of population connectivity and gene flow.

The successful establishment of a bivoltine population in the climatically marginal Hirosaki area also implies rapid local adaptation. This study underscores the usefulness of RAPD markers for detecting fine-scale genetic patterns and tracing invasion histories in introduced insect populations.

## Acknowledgements

We thank members of our laboratory for technical assistance and valuable discussions.

